# Tools for genetic manipulation of the endemic fungal pathogen, *Emergomyces africanus,* and the application of a fluorescent reporter strain in infection models

**DOI:** 10.1101/2025.11.14.688410

**Authors:** Lucian Duvenage, Alisha Chetty, Darren D. Thomson, Elizabeth R. Ballou, Nelesh P. Govender, Chad A. Rappleye, J. Claire Hoving

**Author notes:** Address correspondence to: Lucian Duvenage. (CAR and JCH are co-senior authors).

## Abstract

*Emergomyces africanus* is a thermally dimorphic fungal pathogen endemic to Southern Africa which can cause fatal systemic infections in persons with advanced HIV disease. Its mechanisms of pathogenesis are not well understood. Characterisation of virulence traits in this pathogen requires appropriate molecular tools for genetic manipulation. Molecular technologies developed for the transformation of *H. capsulatum* were adapted for use in *E. africanus. Agrobacterium*-mediated transformation was used to generate a reporter strain expressing green fluorescent protein (GFP). The *E. africanus* GFP reporter strain facilitated the study of yeast interaction with macrophages *in vitro* and allowed the identification of infected phagocyte cell types in the mouse lung by flow cytometry. *E. africanus* could also maintain episomal plasmids with telomere-like sequences, to introduce expression constructs without genome modification. Using this plasmid system, RNA interference constructs were used to knock down the expression of cell wall α(1,3)-glucan by targeting the transcripts of the α-glucan synthase (*AGS1*). An episomal CRISPR/Cas9 system was evaluated for *E. africanus*, which effectively disrupted *GFP* in a reporter strain and enabled the generation of a *URA5* uracil auxotroph. These tools and strains will facilitate future studies to elucidate the mechanisms of pathogenesis of *E. africanus*.

**Importance:** *Emergomyces africanus* is an opportunistic fungal pathogen affecting persons with advanced HIV disease in South Africa. The biology and pathogenesis of *E. africanus* are not well understood, as the importance of the disease caused by this fungus (emergomycosis) has only been recognised in recent years and molecular studies have been impaired by the lack of genetic technologies. In this work, we describe tools and methods for the genetic modification of this pathogen, which will accelerate future studies investigating how the fungus causes disease in the human host. These essential tools include (1) the ability to create fluorescent reporter strains, such as the green fluorescent protein *E. africanus* strain described here, which facilitates tracking the spread of the fungus during infection and enhances microscopy studies, (2) methods for knocking down gene expression in *E. africanus*, and (3) the permanent disruption of genes through CRISPR/Cas9 gene editing.

## Introduction

*Emergomyces africanus*, part of the recently classified genus *Emergomyces* (1), is a fungal pathogen of humans endemic to Southern Africa (2). The fungus can cause life-threatening systemic infection in immunocompromised individuals, which, in South Africa, has exclusively been persons with advanced HIV disease (3). Because this pathogen is less often suspected by clinicians, patients affected by emergomycosis often are diagnosed only at the late stages of infection, characterised by widespread skin lesions. An understanding of its biology and host-pathogen interaction is needed to develop effective diagnostics and treatment strategies for this infection in high-risk populations.

*E. africanus* is a thermally dimorphic fungus. It exists in a saprobic mycelial phase in soil and releases airborne conidia (4), which, when inhaled by the host, transition to a yeast phase triggered by body temperature (5). The mechanisms by which these yeast infect host cells are unknown. The ability to genetically manipulate a fungal pathogen can open new avenues of research, whether by examining the effects of gene deletions or introducing reporter genes for a wide range of applications, such as facilitating the detection of the pathogen in infection models. New tools or protocols must be developed or adapted for non-model fungi such as *E. africanus.* In this study, we used molecular tools originally developed for *Histoplasma capsulatum* and other dimorphic fungi (6–11) and adapted them to the genetic modification of *E. africanus*.

In some fungi, notably the dimorphic fungal pathogens, homologous recombination is inefficient which prevents gene replacement strategies. In these cases, *Agrobacterium-*mediated gene transfer (ATMT) has been employed to introduce genes of interest (9, 11) or fluorescent transcriptional reporters or to create random insertion libraries for phenotypic screening. Co-culture of *A. tumefaciens,* harbouring the shuttle plasmid, with the fungus results in the insertion of a single copy (in the majority of cases) of the T-DNA randomly in the genome (12). This method has been used successfully for several of the thermally dimorphic pathogens, including *Histoplasma capsulatum, Blastomyces dermatitidis, Paracoccidioides brasiliensis* and *Talaromyces marneffei* (as reviewed in (13)). We used ATMT to introduce a GFP transgene into *E. africanus,* using shuttle vectors and a protocol previously developed for the transformation of *H. capsulatum* (14). The *E. africanus* GFP reporter strain facilitated imaging of yeast-macrophage interactions and identification of phagocyte cell types in the lung associated with the yeast in the mouse model of infection (15) by flow cytometry analysis.

Shuttle vectors which replicate in *E. coli* can be maintained by *H. capsulatum* as linear plasmids through the addition of telomere sequences (16–18). These vectors are useful for introducing expression constructs without genome modification. For example, RNA interference (RNAi) constructs can be used to knock down expression of virulence-associated genes (7, 18–22). These plasmids are rapidly lost without selection pressure, which is useful when transient expression is desired, e.g. expression of the genome-modifying enzyme Cas9. Gene deletion/mutation is a powerful tool to study putative virulence-associated traits. CRISPR-Cas9 genome editing has been applied to study these traits in thermally pathogenic fungi, including *Blastomyces dermatitidis* (*23*) and *Histoplasma capsulatum* (*6, 24*). In this study, we applied an episomal plasmid system designed for *H. capsulatum* for CRISPR-Cas9 genome editing (6) in *E. africanus* to delete distinct genes in the genome.

With effective molecular genetic tools in place, and their application optimised for *E. africanus*, its virulence traits and pathogenesis can be investigated in future work. This knowledge could facilitate the development of more specific diagnostic tests for early screening of at-risk patients or inform clinical management of the disease.

## Results and Discussion

### Generation of fluorescent strains using *Agrobacterium*-mediated transformation of a GFP transgene

We used *Agrobacterium tumefaciens-*mediated transformation (ATMT) to transform *E. africanus* using the shuttle vector pAG22, which encodes a GFP transgene. GFP and hygromycin phosphotransferase are expressed under the control of *H. capsulatum* regulatory elements: *TEF1* (translation elongation factor) constitutive promoter and *TUB2 (tubulin)* terminator, and *RPL1B* (large ribosomal subunit protein) promoter and *RPL7* (large ribosomal subunit protein) terminator, respectively (Figure S1). Transformation of *E. africanus* using *A. tumefaciens* LBA1100 or EHA105 strains carrying pAG22 yielded similar numbers of transformants. Following transformation, individual transformants were screened for GFP expression levels, as insertion of the T-DNA in different loci can result in varying levels of GFP expression (12). To identify reporter strains with high GFP expression, transformant colonies were grown in liquid BHI broth and inspected by fluorescence microscopy. Figure 1 shows an example of differences in GFP fluorescence levels between two transformants.

**Figure 1:**
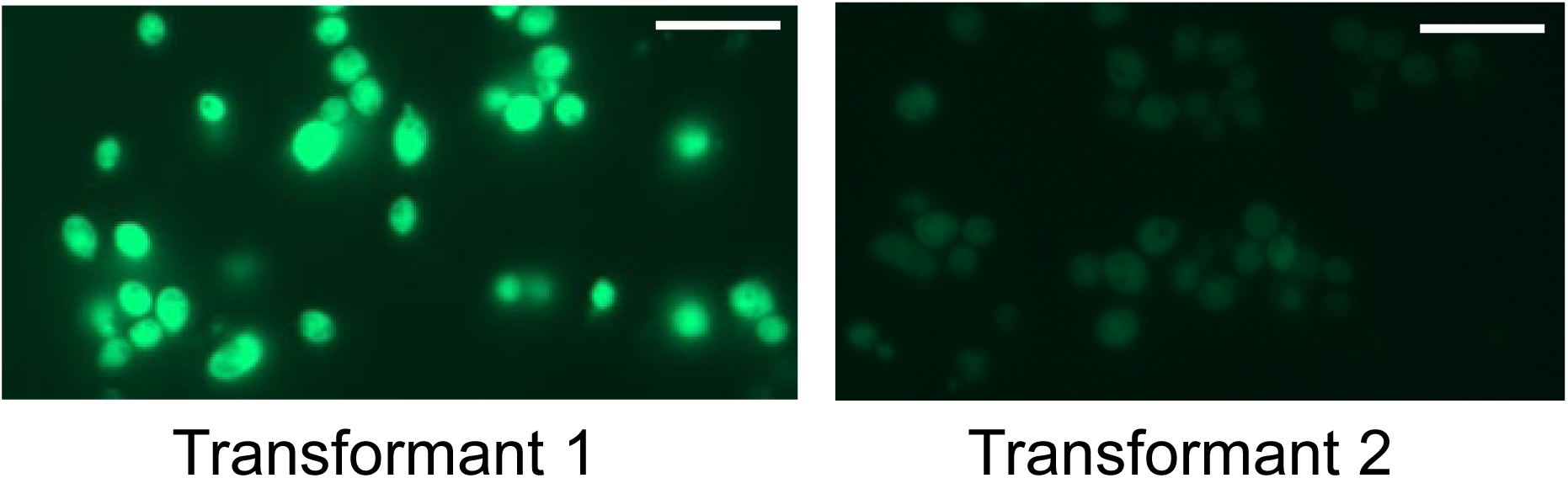
GFP-expressing E. africanus derived through ATMT. E. africanus GFP-expressing transformants were derived by transformation with A. tumefaciens harbouring pAG22. Individual transformant colonies were evaluated by luorescence microscopy, showing variable expression levels of GFP between transformants under constant imaging onditions. Bar=10µm

One pAG22 transformant with high fluorescence was selected for further applications. In preliminary experiments with the mouse model of infection, it was noted that the virulence of laboratory-passaged *E. africanus* (including untransformed wild-type strains) was reduced, as higher doses of yeast were required to replicate pathology compared to previous work. Therefore, all wild-type and control strains used in this study were passaged under the same conditions in the laboratory as would be required to generate respective transformants. An intranasal dose of 1 × 10^7^ yeast cells was then used to infect mice, as this could replicate the pathology seen in previous work (15). The GFP fluorescent strain retained pathogenesis compared to the passage-matched wild-type strain in survival studies (Figure S2).

### Fluorescent E. africanus yeast interaction with macrophages

The GFP-fluorescent yeast facilitated imaging of *E. africanus* interaction with macrophages. J774A.1 macrophages in chambered imaging slides were infected with *E. africanus* yeast (MOI=1) and imaged every 20 minutes. Un-opsonised yeast cells are readily phagocytosed, and a high proportion of yeast cells were intracellular at the start of imaging, one hour following yeast addition. We hypothesise that *E. africanus* may be taken up by macrophages by the same mechanism as *H. capsulatum*, namely, recognition of heat shock protein 60 (Hsp60) on the surface of yeast cells by complement receptor 3 (CD11b/CD18) of macrophages (25), as Hsp60 is highly conserved between the two species. Intracellular yeast could easily be observed by their GFP fluorescence (Figure 2A). Budding of intracellular yeast could be measured in the time-lapse data demonstrating phagocytosed E. africanus yeasts remained viable. The observed budding time of intracellular yeast was not significantly different to that of extracellular yeast in the same proximity (Figure 2B). Yeast cells continued to proliferate intracellularly over time, with some macrophages appearing heavily infected and unable to control yeast proliferation. These data indicate that the intracellular niche is permissive to *E. africanus* replication, as it is for *H. capsulatum.* Previous work showed that bone marrow-derived macrophages are not actively lysed by *E. africanus*, unlike *H. capsulatum*, which may be due to the non-lytic activity of the *E. africanus* Cbp1 homologue (26). These data suggest that, similar to *H. capsulatum* and *P. brasiliensis,* non-activated phagocytes are unable to control *E. africanus,* which may aid in fungal systemic dissemination during infection (27, 28). Further studies using the GFP reporter strain and live cell imaging could provide more information on the dynamics of intracellular replication and macrophage cell death.

**Figure 2:**
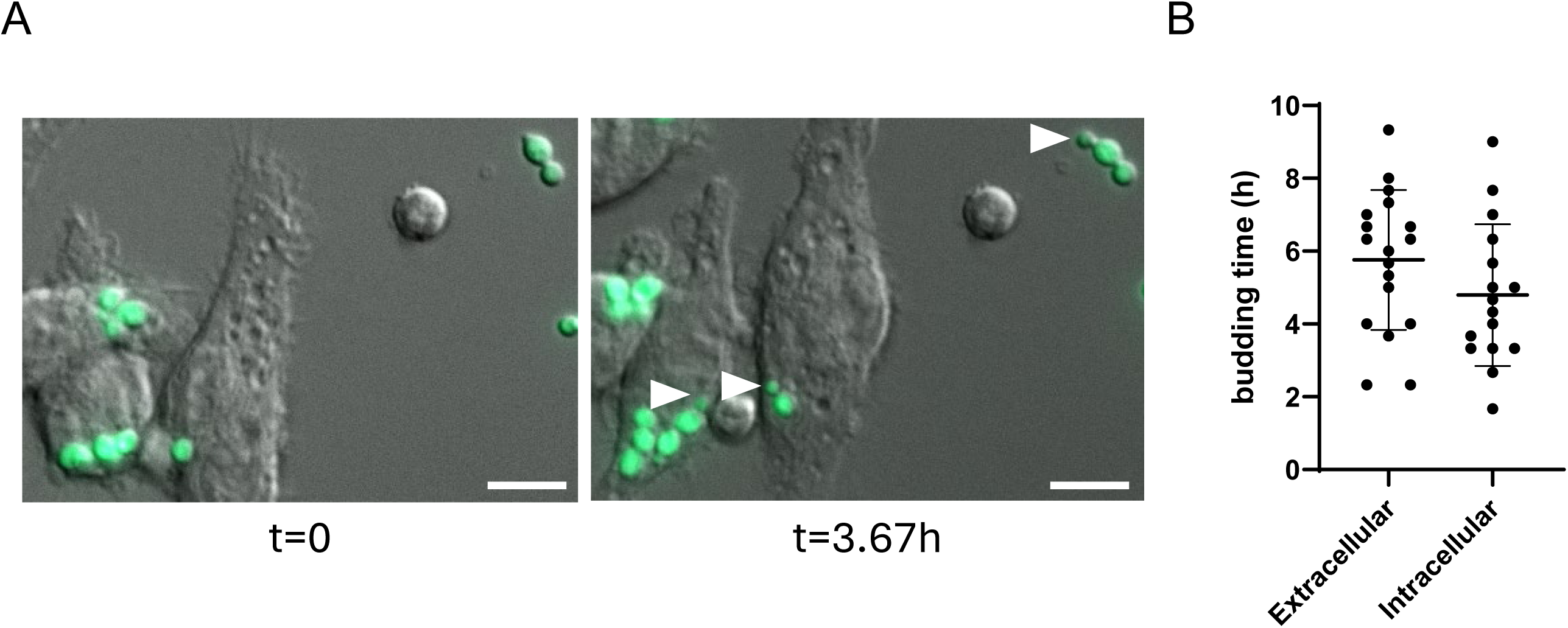
An *E. africanus* GFP fluorescent reporter strain facilitates the study of intracellular yeast growth GFP fluorescent reporter yeast was added to J774.1 macrophage cells in chambered imaging slides. Time-lapse imaging was started 1 hour after yeast addition (t=0), at which point several yeast cells had been internalised (a representative example is shown in A, left). The emergence of new buds (A, right, indicated by arrows) was recorded. Observed budding time was summarised for several fields of view that contained a mix of both extracellular and intracellular yeasts, P>0.05 (mean ± SD). Bar=10µm.

### Identification of infected phagocyte populations in the mouse lung following infection with the GFP reporter strain

We applied the GFP fluorescent *E. africanus* strain to examine its association with phagocytes *in vivo.* Mice were infected intranasally with the fluorescent *E. africanus* strain, and after 7 days, myeloid cell recruitment in the lungs was analysed by flow cytometry. Lungs from uninfected mice were analysed as a control. Gating first on GFP+ cells (i.e., cells with internalised E. africanus) showed that <1% of GFP+ cells were CD45-negative (Figure 3A), suggesting that nearly all phagocytosed *E. africanus* yeasts resided in CD45+ phagocytes. Similar results were reported in a study with an *H. capsulatum* GFP-expressing strain (29), suggesting that both pathogens persist well within this intracellular niche *in vivo*. However, these results only reflect the intracellular fungal burden at the day 7 timepoint, and do not account for fungi that may have been cleared by this time. Most GFP-fluorescent yeasts were associated with neutrophils (CD11b^hi^, Ly6G^hi^) (66.4±17.4% of GFP+ cells; Figure 3B). The remainder of cell types that contained GFP yeast were interstitial macrophages (10.3±5.3%), dendritic cells (MHCII^hi^, CD11c+) (2.1±0.7%) and a very small subset of CD11b-, CD11c- cells (Figure 3B).

**Figure 3:**
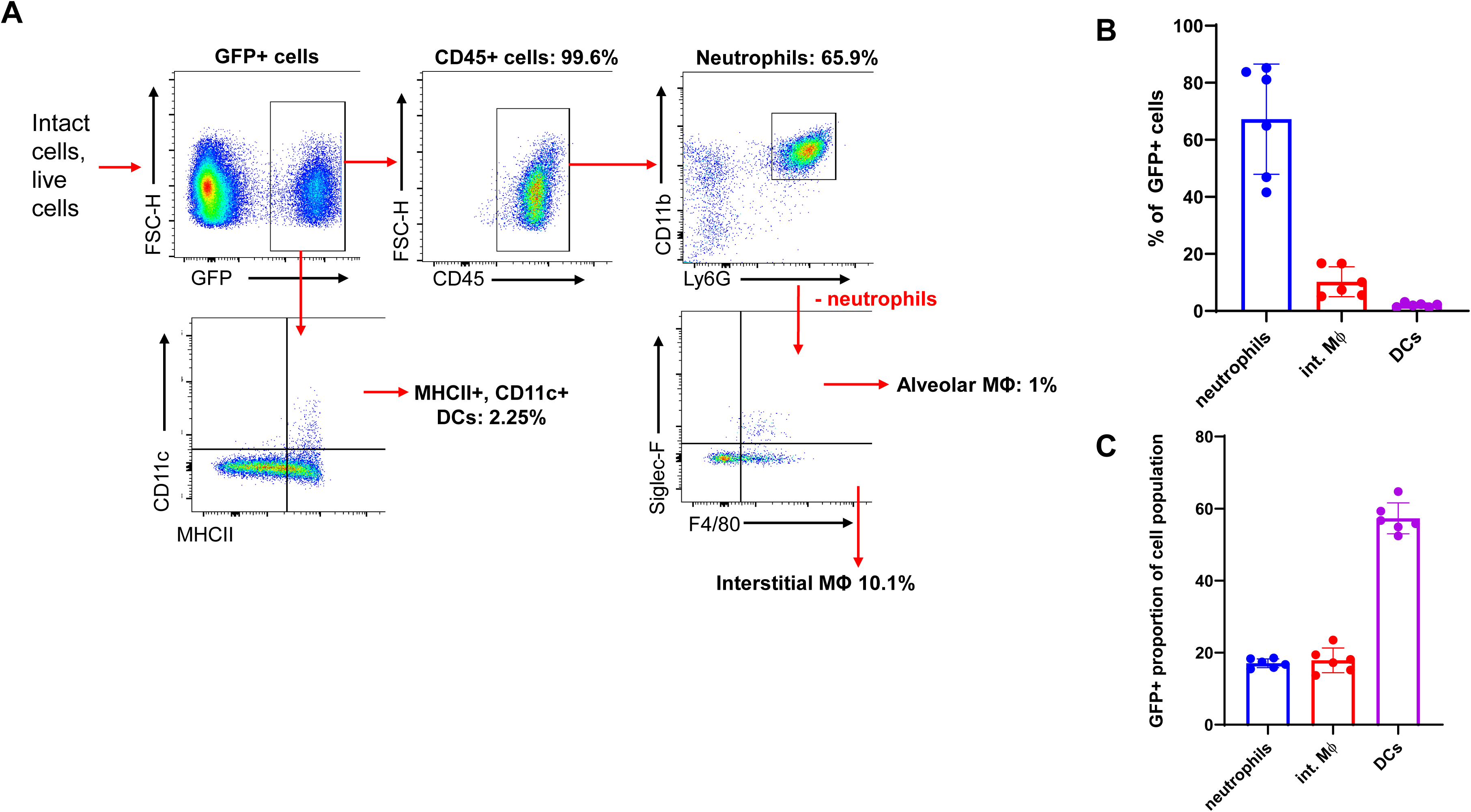
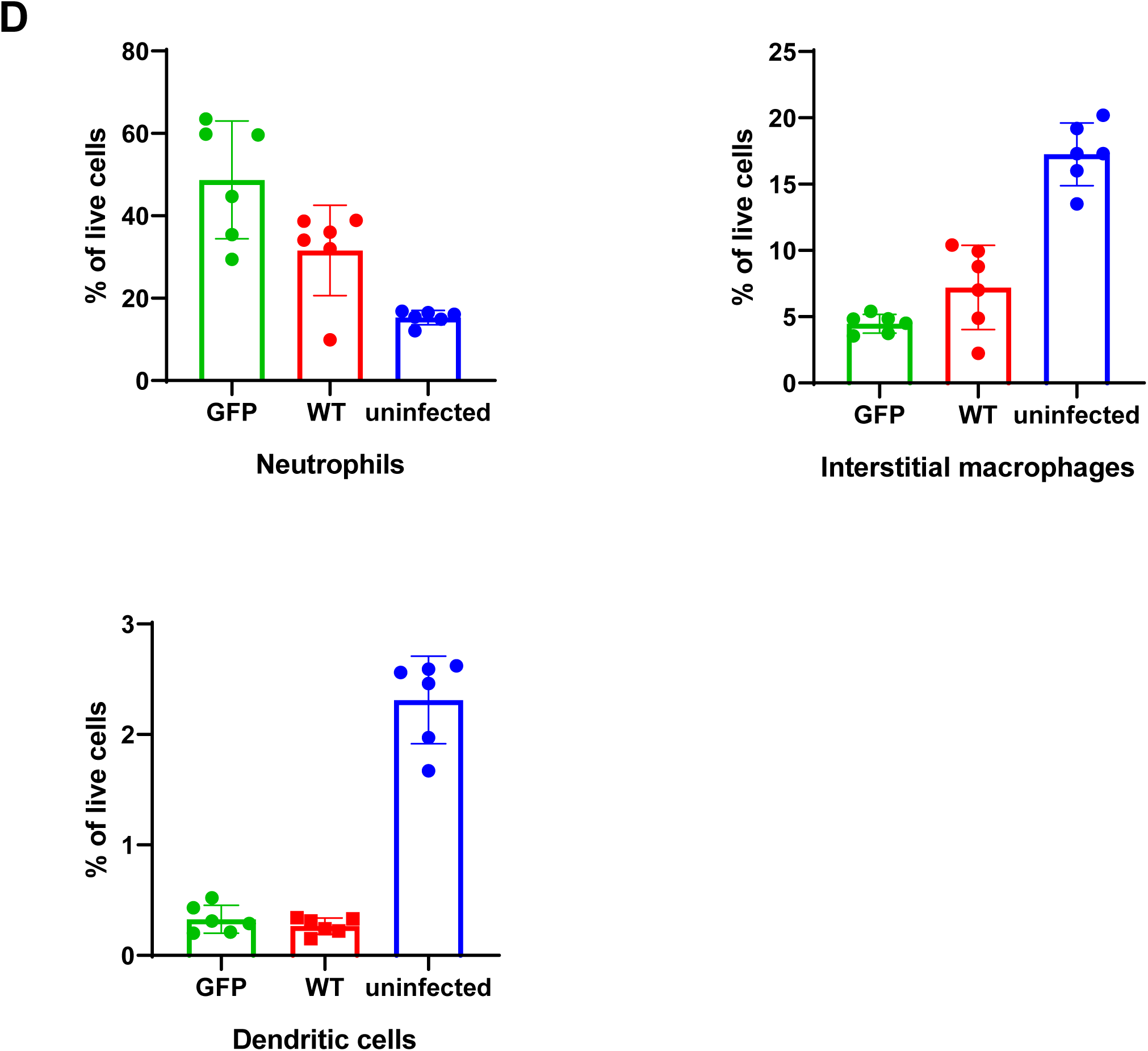
Identification of myeloid cell types associated with GFP-expressing E. africanus in mouse lung at 7 days post-infection. Scatter plots showing the strategy of gating on GFP first (A), which was used to identify myeloid cell types in the mouse lung that were GFP-positive (associated with the GFP reporter yeast), as summarised in (B). (C) Alternate gating on the various cell types first determined the proportions of each cell type associated with GFP yeast. (D) Myeloid cell distribution in the WT-and and GFP strain-infected mouse lung and in uninfected mouse lung. Differences in cell recruitment between WT-and GFP-strain-infected mice were not significant, P>0.05 (mean ± SD). n=6 mice per group, data is representative of two independent experiments.

Alternatively, gating first on all live host cells showed what proportion of total cells were infected with *E. africanus* (GFP+). Of CD45+ cells within the lung, approximately 16.4±4% were infected (GFP+). We then characterised the GFP-fluorescent (i.e., infected) proportion of each myeloid cell type, using a similar gating strategy. The mean proportion of DCs that were infected (GFP-positive) was 57.3±4.3% of DCs detected (MHCII^hi^, CD11c^hi^), with smaller proportions of total lung macrophages and neutrophils infected (Figure 3C). Similar proportions were reported for *H. capsulatum* with the distribution of GFP-fluorescent *H. capsulatum*-infected phagocytes at day 7 post-infection: macrophages: 25.1%, PMNs: 51.2% and DCs: 24.3% for the G217B strain (29). DCs are pivotal in the adaptive immune response to fungi, including *H. capsulatum* (30, 31). The high proportion of DCs infected with *E. africanus* at the 7-day timepoint suggests the initiation of adaptive immunity. The cellular recruitment (of CD45+ cells) to the lung in response to the GFP-fluorescent strain was not significantly different to CD45+ cell populations in animals infected with the non-fluorescent wild-type control (Figure 3D), indicating the GFP-fluorescent E. africanus strain stimulates a similar host immune response. Previous work examined myeloid cell recruitment in *E. africanus*-infected mouse lung at 14 days post-infection, although a lower dose of yeast was used for infection (15). A time course of infection was not performed in the current study; this could be included in future experiments to gain more information on cell recruitment and infected phagocyte proportions over time.

SiglecF+ alveolar macrophages were detected in lungs from uninfected mice (1.5-4% of total cells) but the proportion was reduced in infected mice (0.01-0.16% of total cells). Similarly, the number of interstitial macrophages and dendritic cells was higher in uninfected mice (Figure 3D). This could be due to the proportional increase in neutrophils recruited or apoptosis of leukocytes in the lung, a known feature of infection with *H. capsulatum* and a critical element of protective immunity (32).

### Transformation of *E. africanus* with episomal vectors and reduction of cell wall α(1,3)-glucan by RNA interference

*E. africanus* could be transformed with episomal plasmids based on pCR115 (18) for which linear plasmids with telomeres are maintained without chromosome integration. This was first tested by constructing a modified plasmid with hygromycin B selection (pHygR-GFP). The number of colonies obtained using electroporation of episomal plasmids was typically lower than for ATMT. Hygromycin selection pressure was maintained when working with these transformants, as these plasmids are rapidly lost from the cell population without selection.

To silence production of α(1,3)-glucan in *E. africanus*, we expressed an AGS1-targeting RNA hairpin from an episomal plasmid based on the AGS1-RNAi plasmid pCR115. The RNA hairpin is designed for RNAi-induced knockdown of the *H*. *capsulatum AGS1* gene, with homology to a 678 base pair region of exon 3 of this gene (18). This region is 84% similar overall in *H. capsulatum* and *E. africanus,* with several continuous sections of at least 20 nucleotides or with 100% identity. Therefore, we hypothesised that effective RNAi silencing using the same RNA hairpin could be achieved in *E. africanus,* given that several 21-22 small interfering RNA molecules with complete sequence match are produced following Dicer processing of the long double-stranded RNA. We replaced the GFP in pHygR-GFP with the *AGS1* inverted repeat sequence from pCR115 and used this plasmid (pHygR-HcAGS1-RNAi) to transform *E. africanus.* Yeast transformed with pHygR-HcAGS1-RNAi did not differ in growth rate compared to control yeast, which was transformed with a similar plasmid containing a GFP inverted repeat.

Reduction of cell wall α(1,3)-glucan through RNA interference was assessed by staining the AGS1-RNAi plasmid-transformed yeasts with a monoclonal antibody recognising α(1,3)-glucan. It was noted that not all yeast cells were uniformly stained (Figure 4A) suggesting variable amounts of α(1,3)-glucan. *H. capsulatum* also displays variation in cell wall α(1,3)-glucan levels, as assessed by staining with the same antibody, during broth culture growth (33). Similarly, *E. africanus* cell wall α(1,3)-glucan levels may increase at different time points during culture, or in different growth media. Nevertheless, yeasts transformed with the *AGS1* RNAi plasmid showed an overall lower level of cell wall α(1,3)-glucan staining in the population compared to the control (Figure 4A and 4B). RNAi-induced gene silencing does not typically lead to a complete knockdown of the target, and due to variability in dsRNA expression, a small proportion of cells do not exhibit the full RNAi effect (18). Therefore, some α(1,3)-glucan staining was still observed in *E. africanus* yeasts maintaining the RNAi plasmid.

**Figure 4:**
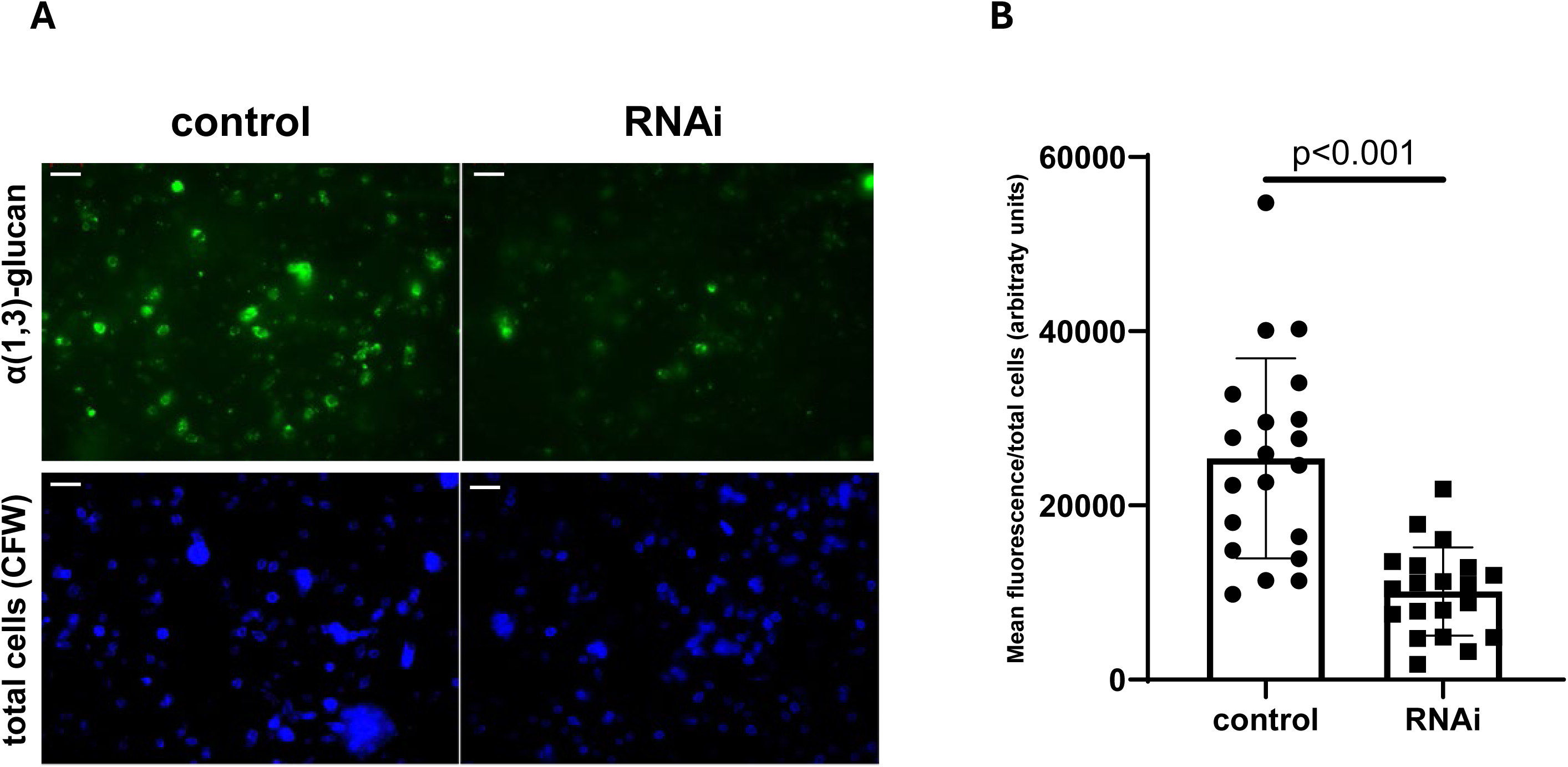
(A) Cell wall staining and fluorescence microscopy of wild-type and AGS1 RNAi strains of E. africanus. E. africanus harbouring the AGS1 RNAi plasmid or the control GFP RNAi plasmid were stained with a monoclonal antibody specific for α(1,3)-glucan and calcofluor white (CFW) and examined by fluorescence microscopy. Imaging settings were kept constant for all samples. Bar=10µm (B) Background-corrected mean fluorescence intensity of microscopy images was measured and normalised by the number of cells per image. Ten images each from two independent experiments were analysed, (mean ± SD) Student’s unpaired t-test, p<0.001.

### Gene mutation in *E. africanus* using an episomal CRISPR/Cas9 system

To test whether CRISPR-Cas9 gene editing could facilitate gene mutation in E. africanus, we transformed a GFP-expressing strain of *E. africanus* with a linear plasmid expressing Cas9 and a GFP-targeting CRISPR gRNA. The GFP-fluorescent *E. africanus* strain was generated through ATMT using pAG21 (GFP-expression and G418 selection). The CRISPR-Cas9 plasmid provided for hygromycin resistance and expressed a gRNA targeting the GFP gene. Hygromycin-resistant transformant colonies were passaged in BHI broth with hygromycin B selection, and the yeasts were examined by fluorescence microscopy for loss of the GFP fluorescence. After successive passages, an increasing proportion of the population was no longer GFP fluorescent (Figure 5A). Sequencing of the GFP gene in the transformant population confirmed a mixed population with indels at the expected Cas9 cut site (Figure 5B) showing efficient CRISPR/Cas9-based genome editing in *E. africanus*.

**Figure 5:**
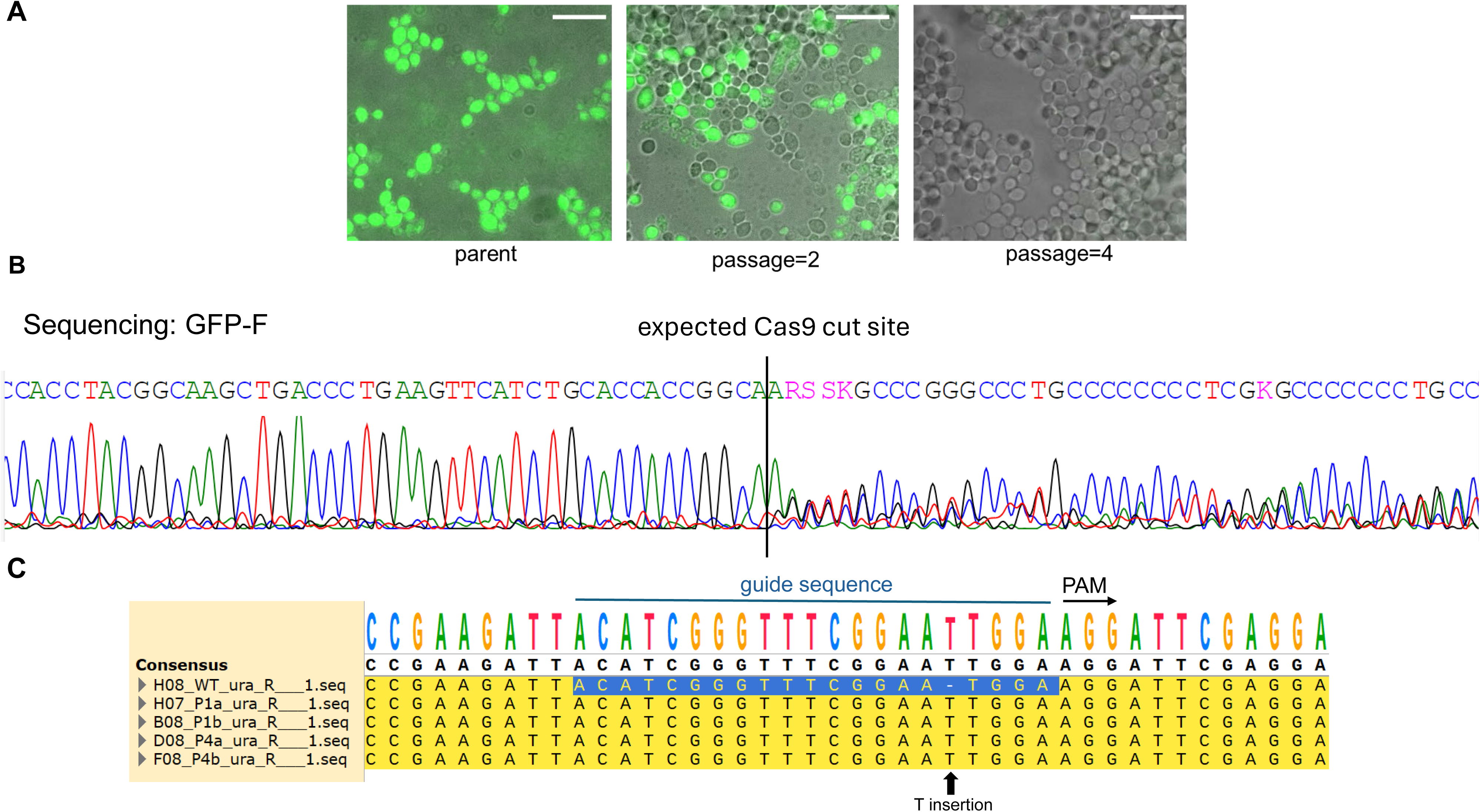
Evaluation of a CRISPR/Cas9 episomal system in E. africanus. (A) Merged fluorescence microscopy and brightfield images showing a decline in the proportion of fluorescent yeast cells with subsequent passages following transformation of GFP-expressing E. africanus with the CRISPR/Cas9 plasmid pSL01-sgGFP. Bar=10µm. (B) Sanger sequencing of a mixed population showed a mixture of signals following the expected Cas9 cut site, indicating the presence of various indels in the population. (C) Sequencing data of individually selected clones, aligned with the WT strain, following CRISPR/Cas9-mediated gene disruption of URA5. A T-insertion was observed in all clones.

Auxotroph strains are useful for genetic manipulations because they facilitate selection of plasmids carrying genes that rescue the auxotrophy. A previous CRISPR/Cas9 workflow used for *H. capsulatum* (6) was used to disrupt *URA5* in *E. africanus* to generate a uracil auxotroph strain. The *E. africanus URA5* homologue (Gene: ACJ72_02173) was identified through a BLAST search. The protein sequence is highly similar to that of *H. capsulatum* (88.1% identity, 93.0% similarity as determined by pairwise alignment using EMBOSS Water (34)). A guide RNA sequence was designed using CRISPOR (https://crispor.gi.ucsc.edu/) (35) to maximise the chance of indel generation in the *URA5* gene while minimising off-target effects. Following transformation of wild-type *E. africanus* with the URA5-targeting CRISPR-Cas9 plasmid, followed by four passages under hygromycin selection, uracil auxotrophs were selected using 5-FOA-resistance screening. Several 5-FOA-resistant clones were identified, and their inability to grow without uracil was validated by failure to grow in media lacking uracil. Sequencing of four clones showed a single nucleotide T insertion in all clones (Figure 5C), causing a frameshift mutation, preventing production of a functional *URA5* gene product. This *E. africanus ura5-*mutant can now be used as the host for *URA5-*based expression vectors or RNAi vectors. These vectors are also stable *in vivo,* as evidenced by continued virulence*; H. capsulatum* uracil auxotrophs are avirulent in mice, which is likely to be the case for *E. africanus* as well (36).

## Conclusions

In this study, we demonstrate that molecular tools developed for genetic modification of *H. capsulatum* could be adapted for use in *E. africanus,* suggesting that the two species use similar regulatory elements for high-level gene expression. Our results include the use of ATMT to transform *E. africanus* with expression plasmids, generating a GFP-fluorescent strain that can be used for tracking infections both in cultured macrophages and *in vivo*. In addition, we provide results showing the efficacy of episomal plasmids designed to induce RNA-interference-based gene knock-down and CRISPR-Cas9-based mutation of target genes. Thus, by adapting technologies, we have put into place tools and methodologies to functionally study genes in E. africanus through expression as well as loss-of-function. Furthermore, we have created an *E. africanus* uracil auxotroph strain enabling the use of URA*5-*based complementation vectors instead of hygromycin selection. Such URA5-based vectors have the advantage of providing for selection *in vivo*, which cannot be done with hygromycin. Together, these tools make *E. africanus* amenable to genetic modification to investigate virulence traits and how they influence the host-pathogen interaction.

## Materials and Methods

### *E. africanus* and *A. tumefaciens* strains and growth conditions

*Emergomyces africanus* (clinical isolate CBS 136260 (4)) was grown in brain-heart infusion broth (BHI) (Cat No. 110493, Merck, USA), at 37 °C, 180 rpm, or maintained on BHI agar plates (Cat. No. 70138, Merck) at 37 °C. *A. tumefaciens* strains LBA1100 and EHA105 was maintained on LB agar plates at 25 °C. Transformation of *Agrobacterium* binary vectors into *A. tumefaciens* was done by electroporation as described in (37) and transformants selected with 100 ug/mL kanamycin.

### Transformation using Agrobacterium tumefaciens

Transformation of *E. africanus* yeast by *Agrobacterium* was done using previously established methods for *H. capsulatum* (14), with the exception that BHI broth or BHI agar plates were used instead of *Histoplasma*-specific growth medium. *Agrobacterium tumefaciens* strains LBA1100 and EHA105 were used to transform *E. africanus* with pAG22 (GFP-expression, hygromycin resistance), yielding similar results. BHI was inoculated with a colony of *E. africanus* from a fresh streak plate, and the culture was grown for three days at 37 °C with shaking. Colonies from a streak plate of *A. tumefaciens* were used to inoculate 5 ml minimal glucose medium (14) for 24 h at 25 °C, then 1 ml of this culture was used to inoculate 5 ml induction medium containing 0.1 mM acetosyringone (Sigma-Aldrich) for a further 16 h growth. *E. africanus* yeast cells were washed with PBS by centrifugation and resuspension, then resuspended in induction medium. The optical density (600 nm) of both *E. africanus* and *A. tumefaciens* suspensions was determined, then combined in 900 µl to give final optical densities of 1.0 for each. Suspensions OD600=1.0 have approximately 5 × 10^8^ cells/ml and 2.5 × 10^8^ cells/ml of *A. tumefaciens* and *E. africanus,* respectively, giving an MOI of 2:1. The co-culture was applied dropwise to sterile Whatman No. 5 qualitative filter papers placed on co-cultivation plates. The co-cultivation plates were incubated at 25 °C for 2 days, then transferred to BHI agar plates containing 300 µg/ml hygromycin B and 200 μM cefotaxime. The plates were incubated at 37 °C for 10-14 days until *E. africanus* transformant colonies appeared. Single colonies were picked and streaked on fresh BHI plates containing 300 µg/ml hygromycin B. From these plates, single colonies were used to inoculate BHI broth to grow liquid cultures of individual transformants. Glycerol stocks (20%) of liquid cultures were preserved at -80 °C.

### Infection of macrophages with *E. africanus* and live-cell imaging

J774A.1 murine macrophage cells (ATCC: TIB-7) were seeded at a density of 2 × 10^5^ cells in 0.3 ml macrophage medium (DMEM high glucose GlutaMAX®, 10% FBS, penicillin/streptomycin [Gibco]) in 8-well imaging slides (ibidi GMBH, Cat. No. 80826) 24 h prior to the addition of *E. africanus* yeast. *E. africanus* yeast was grown for 3 days in BHI broth, then washed with PBS and counted using a haemocytometer. 2 × 10^5^ yeast cells in 300 µl macrophage media were added to macrophages in imaging slides to achieve a ratio of one yeast cell per macrophage (MOI=1). The slides were then set up for time-lapse imaging under the conditions of 37 °C, 5% CO2. Imaging was started 1 hour post-yeast addition. Imaging was done using a Zeiss AxioObserver fluorescence microscope at 40X magnification. Analysis of observed budding time was done by examining yeast cells that were intracellular or extracellular, at t=0, with no visible buds.

### Mouse model of infection and flow cytometry

To examine the responses to *E. africanus* in the mouse lung, three groups of mice were analysed (n=6 per group): (1) uninfected mice, (2) mice infected with the wild-type clinical isolate and (3) mice infected with the GFP fluorescent *E. africanus* strain. Eight- to twelve-week-old male CL57/B6 mice were obtained from the University of Cape Town Animal Research Specific Pathogen Free Facility. *E. africanus* wild-type and GFP fluorescent strains were grown in BHI broth for 3 days at 37 °C until OD600 reached approximately 1.0. Yeast cells were washed twice with PBS before counting using a haemocytometer. Mice were anaesthetised with ketamine and xylazine (80 mg/kg and 16 mg/kg, respectively), then infected intranasally with 50 µl of yeast cell suspension containing 1 × 10^7^ yeast cells. Animal welfare was monitored daily. After 7 days, mice were euthanised by halothane inhalation and the lungs were removed and stored in DMEM + 5% FBS on ice. Lungs were digested in digestion buffer (15) containing 50 units/ml Collagenase Type I and 13 μg/ml DNAseI for 1h at 37 °C. The digested lungs were passed through a 70 µm cell strainer using a sterile syringe plunger. This process was repeated using a 40 µm cell strainer. Red blood cells were lysed using 1x RBC lysis buffer (eBioscience), then remaining cells collected by centrifugation and resuspension in DMEM + 5% FBS, then counted using haemocytometer. 2 × 10^6^ cells per well were added to a V-bottom 96-well plate. Myeloid cells were stained in MACS buffer (0.5% BSA, 2 mM EDTA in PBS) using the antibodies in Table 3, together with 1% Fc block and 2% heat-inactivated rat serum, for 20 min at 4 °C in the dark, then washed twice with MACS buffer. 7-AAD (Biolegend) was used as the cell viability stain. Single-stained and unstained controls were used to compensate for spectral overlap. Stained cells were analysed by flow cytometry using an LSR Fortessa™ (BD Immunocytometry Systems, San Jose, CA, USA) and BD FACS Diva software (v6.0). Flowjo software (v10.0.7) (Treestar, Ashland, OR, USA) was used for post-acquisition analysis cell population determination. The gating strategy used is shown in Figure 3A.

### Ethics statement

All work with mice was done with ethical clearance from the University of Cape Town Animal Research Ethics Committee, reference numbers 020/017 and 024/014.

### Statistical analysis and software

Values are reported as means ± standard deviation. Differences between groups were determined using a two-tailed Student’s t-test or one-way ANOVA using GraphPad Prism v10.6.1. Plasmid map generation and alignments of DNA sequence data were done using SnapGene v8.02. ImageJ 1.54p was used for image analysis (mean fluorescence intensities and total cell counting).

### Plasmid construction

Plasmids pAG21, pAG22, pCR115 (18), pSL01 and pCR745 (6) were provided by CAR. Isolation of plasmid DNA, restriction enzyme cloning and PCR/gel purification was done using standard methods, and plasmids were maintained in *E. coli* DH5α. Restriction enzymes and ligation kits were purchased from New England Biolabs. An episomal GFP expression plasmid (pHygR-GFP) was constructed by digestion of pAG22 with EcoRI and HindIII, and ligating the fragment containing the GFP- and *hph* expression cassettes into the similarly-digested pCR115 vector backbone. To construct pHyg-HcAGS1-RNAi (AGS1 inverted repeat) used for *AGS1* RNAi, the *GFP* sequence from plasmid pHygR-GFP was exchanged with the *AGS1* inverted repeat sequence from pCR115 by digestion with AscI and SpeI. A control *GFP* RNAi plasmid (pHygR-GFP-RNAi) was constructed by inserting an inverted repeat of *GFP* into pHygR-GFP by PCR with GFP-SpeI-F and XhoI-GFP-R primers and restriction enzyme cloning. For CRISPR/Cas9-mediated disruption of GFP, plasmid pCR115-sgGFP was created: a portion of plasmid pCR745 (6) containing the guide RNA targeting GFP, was digested with SwaI and AvrII and ligated into similarly digested pSL01 (6), resulting in plasmid pSL01-sgGFP. For CRISPR/Cas9-medited disruption of *URA5,* a synthetic DNA sequence containing the guide RNA sequence, hammerhead and HDV ribozyme sequences, and part of the tracr RNA sequence, including 15 bp up- and downstream-homology regions was designed for cloning into pSL01 as described in (6). The guide RNA sequence was designed using CRISPOR (https://crispor.gi.ucsc.edu/) (35) to maximise the chance of indels while preventing off-target effects. This sequence was synthesised by Integrated DNA Technologies (Leuven, Belgium). The synthetic DNA sequence was cloned into SwaI-digested pSL01 and transformed into competent *E. coli* cells using In-Fusion® HD Cloning Kit (Takara Bio) using the manufacturer’s recommendations. Table 1 summarises the plasmids used in this study.

**Table 1:** Plasmids used in the study.

| Plasmid | Application | Selection |
| --- | --- | --- |
| pAG22 | ATMT | Hygromycin B (300 µg/ml) |
| pAG21 | ATMT | G418 (1 mg/ml) |
| pCR115 | AGS1 RNAi | [Uracil auxotrophy] |
| pHygR-GFP | GFP episomal plasmid | Hygromycin B (300 µg/ml) |
| pHygR-GFP-RNAi | GFP inverted repeat plasmid (RNAi control) | Hygromycin B (300 µg/ml) |
| pHygR-HcAGS1-RNAi | AGS1 RNAi | Hygromycin B (300 µg/ml) |
| pCR745 | CRISPR/Cas9 | [Uracil auxotrophy] |
| pSL01 | CRISPR/Cas9 | Hygromycin B (300 µg/ml) |
| pSL01-sgGFP | CRISPR/Cas9 | Hygromycin B (300 µg/ml) |
| pSL01-sgURA | CRISPR/Cas9 | Hygromycin B (300 µg/ml) |

### Transformation of yeast with episomal plasmids by electroporation

Transformation of *E. africanus* with plasmids by electroporation of was based on previously established methods for *H. capsulatum* (17). Briefly, *E. africanus* yeasts were grown in 5 ml BHI broth for 3 days until OD600 reached approximately 1.0. Yeast cells were collected by centrifugation and resuspended in 2 ml 10% mannitol. Plasmids were linearised by digestion with PmeI to expose the telomeres, then column-purified and eluted with nuclease-free water. Three hundred microlitres of *E. africanus* yeast suspension (1 × 10^7^ yeast cells) was mixed with 200 ng purified plasmid in a chilled electroporation cuvette (0.2 cm gap), then electroporated in a single pulse using a Bio-Rad Gene Pulser® II system with the following settings: 1.25 kV, 50 mF, 600 Ω, which gave a time constant of 8-12 ms. Following electroporation, 500 µl of BHI broth was mixed with cells and this cell suspension was applied dropwise to sterile Whatman No. 5 qualitative filter papers placed on BHI agar plates with no selection. These plates were incubated for 48 h at 37 °C to allow for recovery of transformants, before transferring the membranes to BHI agar plates containing 300 µg/ml hygromycin B. The plates were incubated at 37 °C for 10-14 days until transformant colonies appeared. Single colonies were picked and streaked onto fresh BHI plates (300 µg/ml hygromycin B). Selection with hygromycin B (300 µg/ml) was maintained for growing liquid cultures, after which -80 °C glycerol stocks were prepared for long-term storage of strains, as described above.

### Cell wall alpha(1,3)-glucan staining of RNAi strains

Yeasts harbouring the *AGS1* RNAi plasmid pHygR-HcAGS1-RNAi or the control plasmid pHygR-GFP-RNAi were grown for 3 days in BHI broth with 300 µg/ml hygromycin B until cell density reached an OD600 of 1.0. Cells from 300 µl of culture were collected by centrifugation, then washed twice with PBS, and fixed in 4% paraformaldehyde overnight at 4 °C. Cells were washed twice with PBS, then blocked with 1% BSA, PBS for 1 h. Cells were then stained with 1:150 mouse IgM MOPC-104E antibody (M5170, Sigma-Aldrich) in 1% BSA, PBS for 2 h at ambient temperature. The cells were washed twice by centrifugation and resuspended in PBS and stained with 1:400 anti-mouse IgM secondary antibody conjugated to AlexaFluor488 (Invitrogen, A-21042) for 1 h at ambient temperature. To stain total yeast cells, calcofluor white was added at a final concentration of 100 µg/ml and incubated for 15 min at ambient temperature in the dark. Cells were washed twice with PBS and resuspended in 20 µl PBS. A 5µl sample of stained cells was spotted onto a microscope slide, allowed to air-dry and mounted with ProLong™ Glass Antifade Mountant (P36984, Invitrogen). The slides were examined using a Zeiss Axiovert 200M fluorescence microscope with a monochrome Zeiss AxioCam HRm camera at 100x magnification. Exposure times were kept constant for all samples when capturing images for mean fluorescence analysis. Fluorescence of the population was determined by measuring the background-corrected mean grey values for 10 fields of view each using ImageJ, for two pooled independent experiments, then dividing this value by the number of total cells (as determined by calcofluor white staining).

### Targeted mutagenesis using CRISPR/Cas9-plasmids

A GFP-expressing strain of *E. africanus* was generated through ATMT using pAG21 (equivalent to pAG22 but with a G418 resistance marker instead of hygromycin B resistance). The GFP-fluorescent strain was subsequently transformed with PmeI-linearised plasmid pSL01-sgGFP by electroporation. Hygromycin-resistant transformant colonies were picked and grown in BHI broth for successive passages (1:50 dilution), maintaining hygromycin selection every 3 days. Samples of these cultures were examined by fluorescence microscopy for GFP expression. For DNA sequencing, yeast cells from 1 ml of culture was collected by centrifugation, lysed with 200 µl of lysis buffer (2% Triton X-100, 20 mM EDTA) and beating (2 min) with approximately 100 µl of glass beads (0.5 µm diameter). 100 µL of phenol and 100 µl chloroform-isoamyl alcohol (25:24:1) were added, and the phases were separated by centrifugation. DNA was recovered by transferring 50 μl of the aqueous (upper) phase into 200 μl distilled water. This crude DNA-containing solution was used as the template (2% vol/vol) for PCR using Taq polymerase and primers flanking the expected gene editing sites: GFP-SpeI-F and GFP-Xho-R, or Ura seq-F and Ura seq-R.

For mutation of *URA5* to create a uracil auxotroph, a guide RNA sequence targeting URA5 was designed using CRISPOR (https://crispor.gi.ucsc.edu/) (35). A DNA construct containing the crRNA, hammerhead- and HDV ribozyme sequences, and part of the tracrRNA sequence, with 15 bp flanking homology arms, was synthesised as described in (6) for cloning into plasmid pSL01, by Integrated DNA Technologies (Leuven, Belgium). The sequence of this construct is shown in Table 2. Cloning of this synthetic DNA into SwaI-digested plasmid pSL01 was done through InFusion Cloning (Takara Biotec) as per the manufacturer’s recommendations. Constructs were verified through Sanger sequencing. *E. africanus* was transformed with this plasmid by electroporation as described above, followed by selection with hygromycin B. After four passages in liquid culture, cultures were streaked onto BHI plates supplemented with 1mg/ml 5-Fluoroorotic Acid (5-FOA, R0811, ThermoFisher) and 100 µg/ml uracil to isolate uracil auxotrophs. Uracil auxotrophy was confirmed by the inability to grow in DMEM/F12 media (11320033, Gibco), which lacks uracil, compared to growth in parallel in DMEM/F12 media supplemented with 100 µg/ml uracil. Independent clones were picked for Sanger sequencing.

**Table 2:**
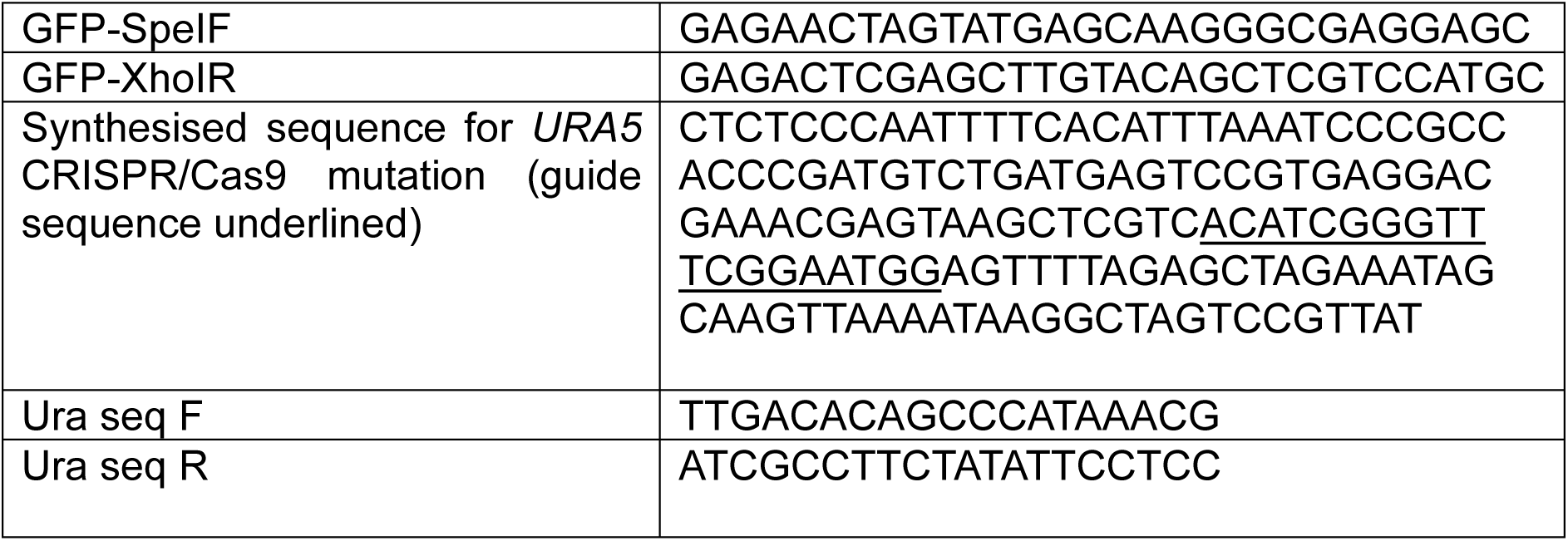
Primers and oligonucleotides.

**Table 3:** Flow cytometry antibodies.

| Target (Clone) | Conjugated fluorophore | Manufacturer (catalogue number) | Dilution |
| --- | --- | --- | --- |
| CD45 (30-F11) | BV510 | BD Biosciences (567800) | 1:100 |
| CD11b (M1/70) | BV421 | BD Biosciences (566313) | 1:100 |
| CD11c (HL3) | Alexa Fluor 700 | BD Biosciences (560583) | 1:100 |
| Ly-6G (1A8) | APC | BD Biosciences (560599) | 1:100 |
| F4/80 (BM8) | PE/Cy7 | Biolegend (123113) | 1:100 |
| Siglec-F (E50-2440) | APC/Cy7 | BD Biosciences (565527) | 1:50 |
| I-A/I-E (MHCII) (M5/114.15.2) | PE/Cy5 | Biolegend (107611) | 1:100 |

---

**Figure S1:**
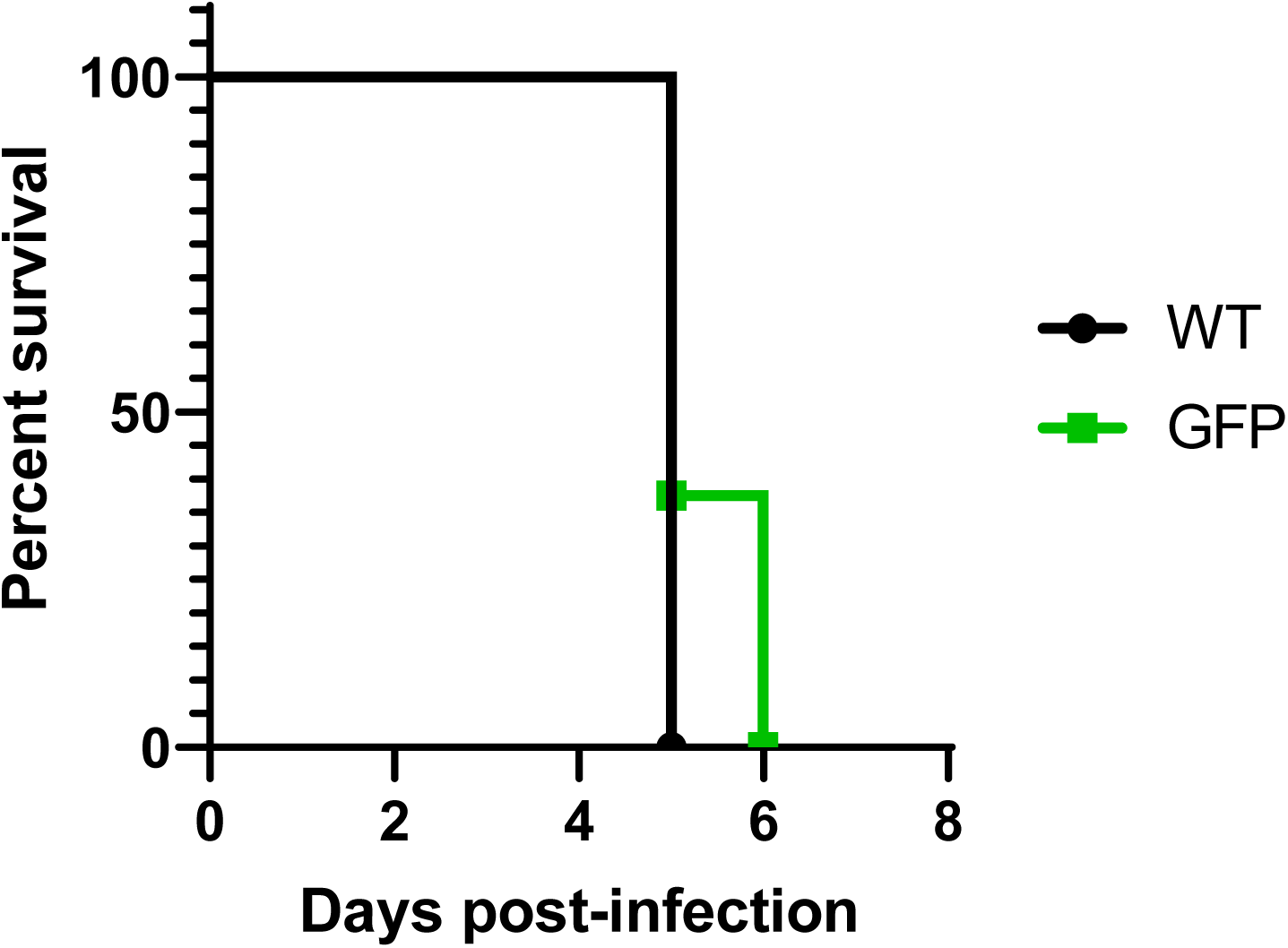
Survival of mice infected with wild-type *E. africanus* and the GFP reporter strain. Mice (n=8 per group) were infected intranasally with a lethal dose of passage-matched wild-type or GFP reporter *E. africanus* yeast cells (2 × 10^7^). Mice were weighed daily and euthanised at the humane endpoint (20% weight loss). The two survival curves are not significantly different, as determined by the log-rank test.

**Figure S2:**
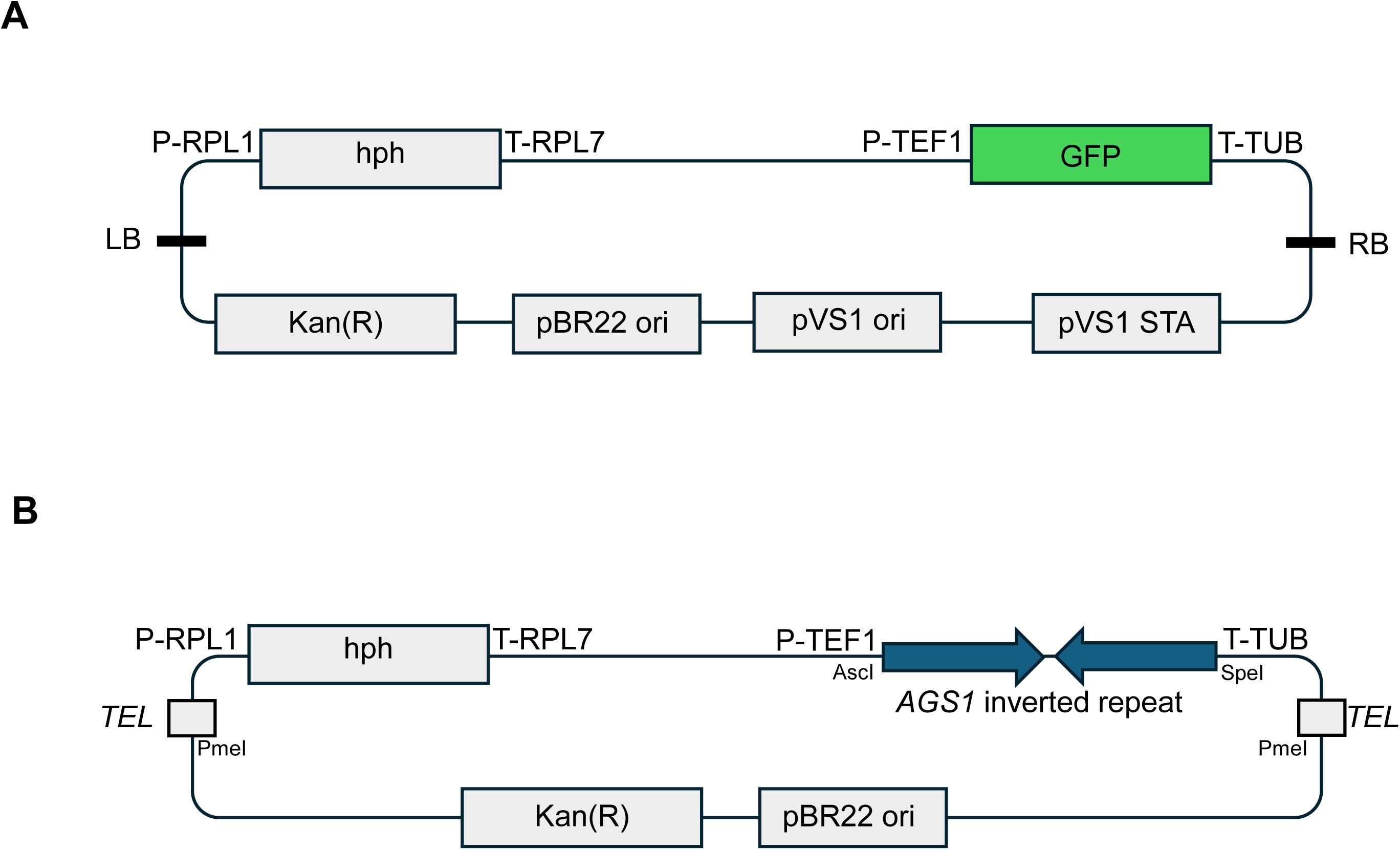
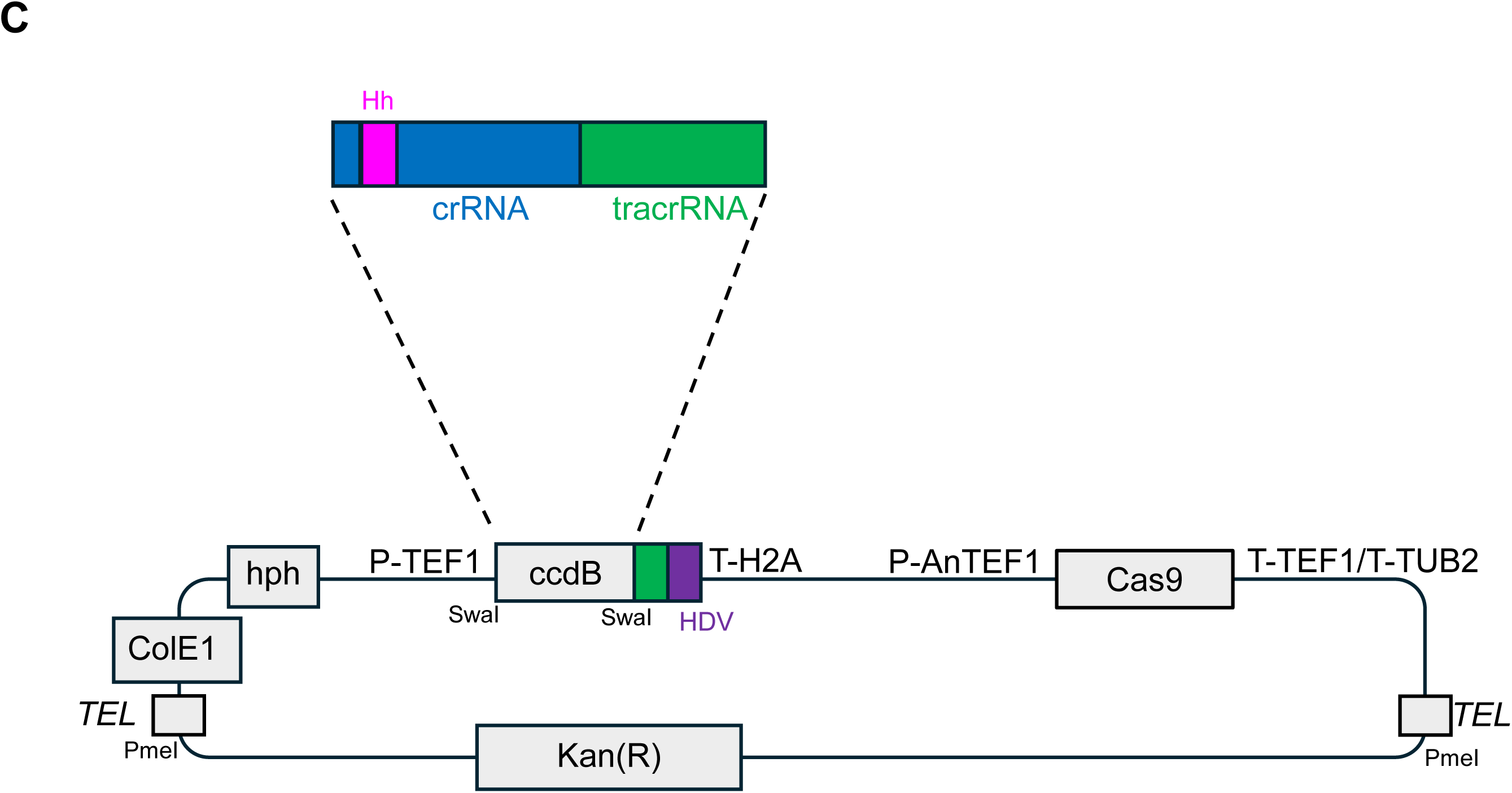
Schematics of plasmids used in this study with selected features. A: *Agrobacterium* shuttle vector pAG22 contains origins of replication for *E. coli* and *A. tumefaciens,* kanamycin resistance marker, a hygromycin resistance marker under the control of *RPL1* promoter and *RPL7* terminators, and GFP under the control of *TEF1* promoter and *TUB* terminator. LB/RB designate the left- and right borders of the T-DNA. All promoter and terminator sequences are derived from *Histoplasma capsulatum* G217B sequences. B: Telomeric plasmid with a hygromycin resistance marker used for RNAi. A linearised plasmid with telomere ends (TEL) was generated by PmeI digestion before transforming *E. africanus*. The *AGS1* inverted repeat sequence was derived from the *H. capsulatum* sequence as described in (18). C: Episomal CRISPR/Cas9 plasmid pSL01, as described in (6), was used to induce frameshift mutations in *E. africanus*. The plasmid was digested with SwaI, and the ccdB marker was replaced by an oligonucleotide containing the *URA5* crRNA sequence with upstream inverted repeat, Hh ribozyme sequence, and part of the tracrRNA sequence, resulting in the expression of a chimeric guide RNA molecule (gRNA), processed by Hh and HDV ribozyme cleavage. The gRNA expression is under the control of the *TEF1* promoter *H2A* terminator derived from *H. capsulatum.* Cas9 expression is driven by the *Aspergillus nidulans TEF1* promoter. The plasmid is linearised by PmeI digestion before transformation of *E. africanus* by electroporation.

