## Supplementary Figures S1 and S2 for "Tools for genetic manipulation of the endemic fungal pathogen, *Emergomyces africanus,* and the application of a fluorescent reporter strain in infection models"

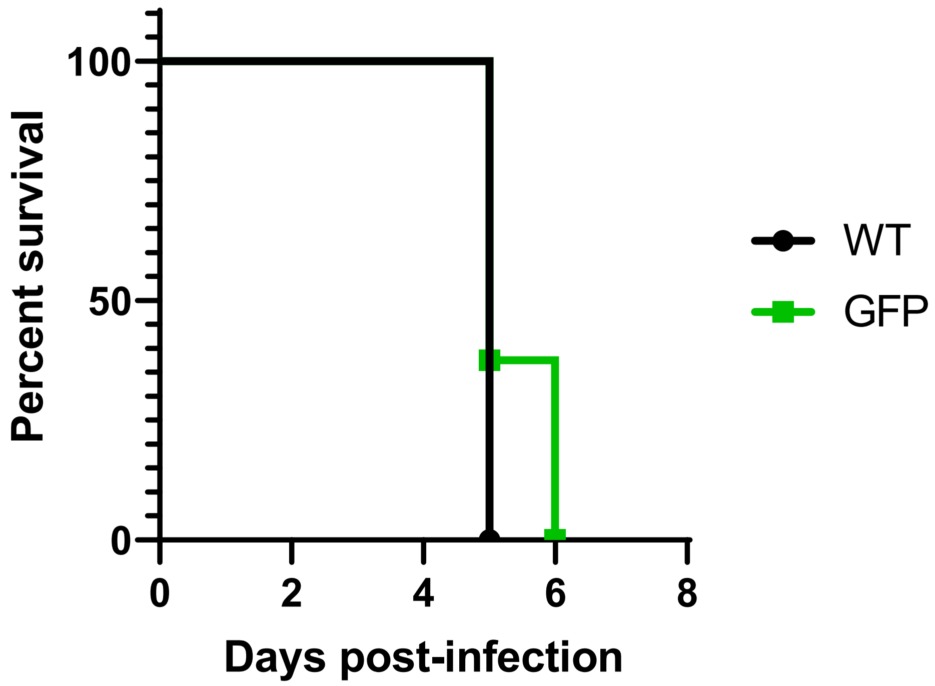

100
50
0
Percent survival
0
2
4
6
8
Days post-infection
WT
GFP

**Figure S1: Survival of mice infected with wild-type *E. africanus* and the GFP reporter strain**

Mice (n=8 per group) were infected intranasally with a lethal dose of passage-matched wild-type or GFP reporter *E. africanus* yeast cells ($2 \times 10^7$). Mice were weighed daily and euthanised at the humane endpoint (20% weight loss). The two survival curves are not significantly different, as determined by the log-rank test.

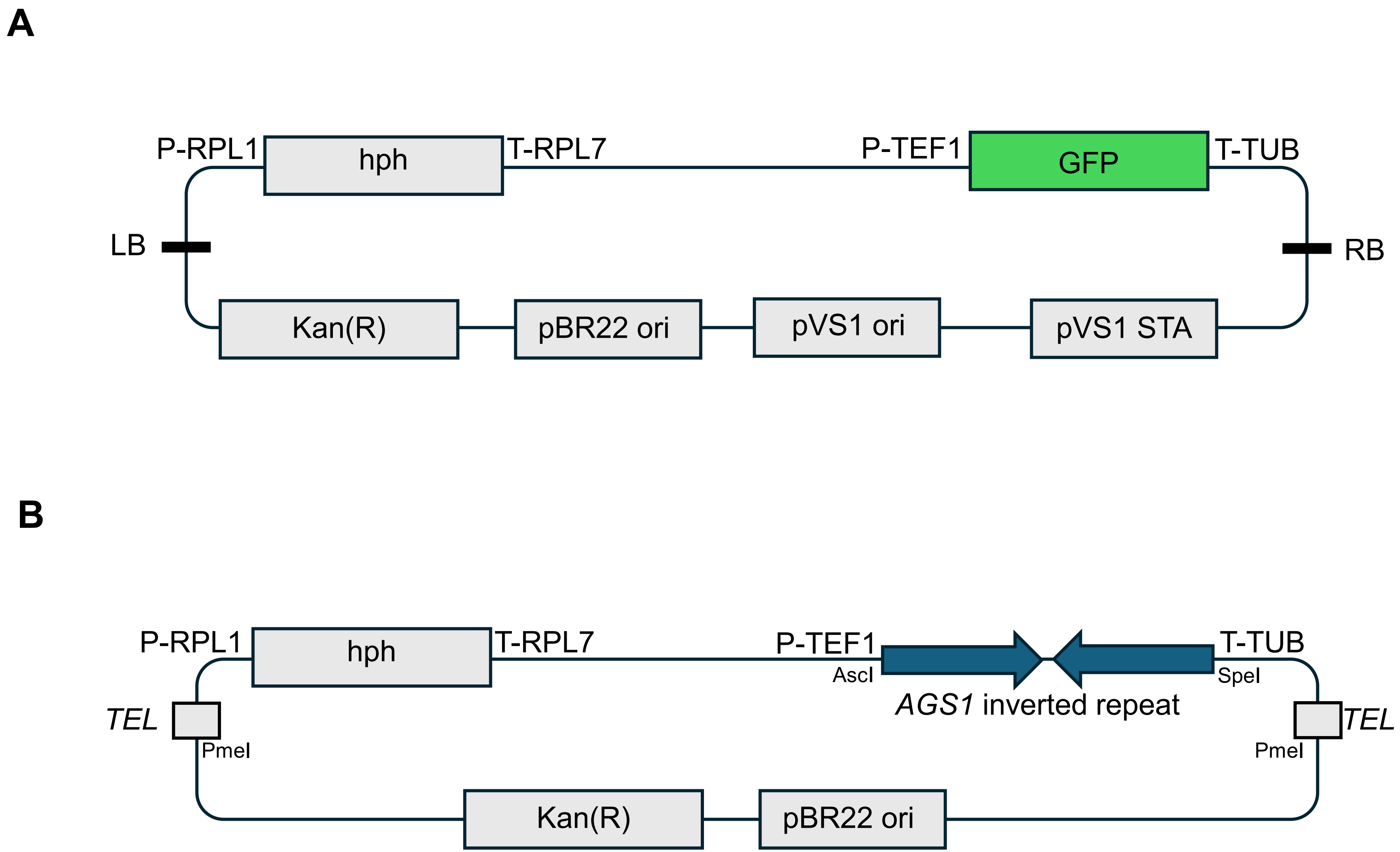
A
P-RPL1
hph
T-RPL7
P-TEF1
GFP
T-TUB
LB
RB
Kan(R)
pBR22 ori
pVS1 ori
pVS1 STA
B
P-RPL1
hph
T-RPL7
P-TEF1
AscI
*AGS1* inverted repeat
T-TUB
SpeI
*TEL*
PmeI
*TEL*
PmeI
Kan(R)
pBR22 ori

C

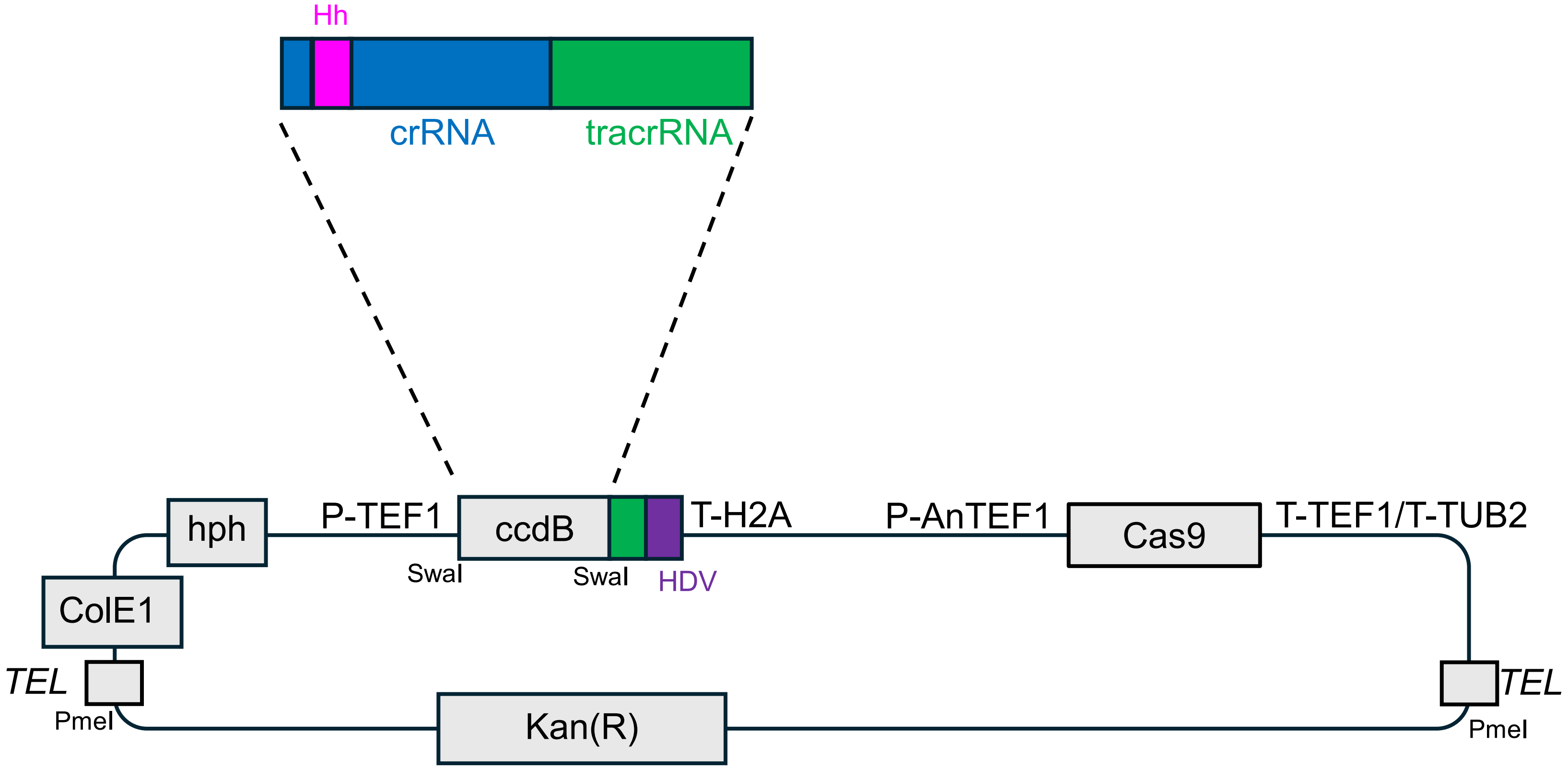
Hh
crRNA
tracrRNA
hph
P-TEF1
ccdB
T-H2A
P-AnTEF1
Cas9
T-TEF1/T-TUB2
SwaI
SwaI
HDV
ColE1
TEL
PmeI
Kan(R)
TEL
PmeI

**Figure S2: Schematics of plasmids used in this study with selected features**

A: *Agrobacterium* shuttle vector pAG22 contains origins of replication for *E. coli* and *A. tumefaciens,* kanamycin resistance marker, a hygromycin resistance marker under the control of *RPL1* promoter and *RPL7* terminators, and GFP under the control of *TEF1* promoter and *TUB* terminator. LB/RB designate the left- and right borders of the T-DNA. All promoter and terminator sequences are derived from *Histoplasma capsulatum* G217B sequences.

B: Telomeric plasmid with a hygromycin resistance marker used for RNAi. A linearised plasmid with telomere ends (TEL) was generated by PmeI digestion before transforming *E. africanus*. The *AGS1* inverted repeat sequence was derived from the *H. capsulatum* sequence as described in (18).

C: Episomal CRISPR/Cas9 plasmid pSL01, as described in (6), was used to induce frameshift mutations in *E. africanus*. The plasmid was digested with SwaI, and the ccdB marker was replaced by an oligonucleotide containing the *URA5* crRNA sequence with upstream inverted repeat, Hh ribozyme sequence, and part of the tracrRNA sequence, resulting in the expression of a chimeric guide RNA molecule (gRNA), processed by Hh and HDV ribozyme cleavage. The gRNA expression is under the control of the *TEF1* promoter *H2A* terminator derived from *H. capsulatum.* Cas9 expression is driven by the *Aspergillus nidulans TEF1* promoter. The plasmid is linearised by PmeI digestion before transformation of *E. africanus* by electroporation.
